# Population dynamics of mutualism and intraspecific density dependence: how *θ*-logistic density dependence affects mutualistic positive feedback

**DOI:** 10.1101/108175

**Authors:** Christopher M. Moore, Samantha A. Catella, Karen C. Abbott

## Abstract

Mutualism describes the biological phenomenon where two or more species are reciprocally beneficial, regardless of their ecological intimacy or evolutionary history. Classic theory shows that mutualistic benefit must be relatively weak, or else it overpowers the stabilizing influence of intraspecific competition and leads to unrealistic, unbounded population growth. Interestingly, the conclusion that strong positive interactions lead to runaway population growth is strongly grounded in the behavior of a single model. This model ― the Lotka-Volterra competition model with a sign change to generate mutualism rather than competition between species ― assumes logistic growth of each species plus a linear interaction term to represent the mutualism. While it is commonly held that the linear interaction term is to blame for the model’s unrealistic behavior, we show here that a linear mutualism added to a *θ*-logistic model of population growth can prevent unbounded growth. We find that when density dependence is decelerating, the benefit of mutualism at equilibrium is greater than when density dependence is accelerating. Although there is a greater benefit, however, decelerating density dependence tends to destabilize populations whereas accelerating density dependence is always stable. We interpret these findings tentatively, but with promise for the understanding of the population ecology of mutualism by generating several predictions relating growth rates of mutualist populations and the strength of mutualistic interaction.

## 1. Introduction

Mutualistic interactions describe the ecology of two or more species that reciprocally increase each other’s fitness (Bronstein, 2015). These interactions are arguably the most common type of ecological interaction, and they have profoundly shaped biodiversity as we understand it. Examples include mutualisms bet5 ween mycorrhizae and plants (van der Heijden et al., 2015), coral and zooxanthellae (Baker, 2003), plants and pollinators (Willmer, 2011), tending ants and aphids or Lepidoptera larvae (Rico-Gray and Oliveira, 2007; Stadler and Dixon, 2008), plants and seed-dispersing animals (Howe and Smallwood, 1982; Levey et al., 2002), lichens (fungi and algae) (Brodo et al., 2001), and plants and rhizobia (Sprent et al., 1987; Kiers et al., 2003). Despite mutualism’s obvious importance, it was not until the latter part of the 20^th^ century that the natural history of mutualism underwent rigorous ecological study, the conceptual framework for mutualism was laid, and mutualism was no longer confounded with the concept of symbiosis. Thus, by the time mutualism was fully introduced to the larger ecological community, theoretical ecology had been developing in its absence for decades. This resulted in the paucity of theory for mutualisms still very much visible today.

Gause and Witt (1935) first used the Lotka-Volterra model of interspecific competition to investigate the case of facultative “mutual aid” between two species by reversing the sign of the linear competition term from negative to positive. They noted that with enough “mutual aid” the zero-growth isoclines no longer cross to give a positive equilibrium point and species grow exponentially without bound—a biologically unrealistic scenario. More specifically, they found that if the product of the strength of mutualism between the two species is ≥ the product of the strength of intraspecific competition for each species, then the positive feedback of mutualism would overpower the negative feedback of intraspecific competition, resulting in unbounded growth. Following this pioneering study, no development of theory around mutualism would happen for over 30 years and ecologists were left lacking a basic theoretical explanation for what stabilizes mutualism in nature.

A key feature of the Lotka-Volterra model is its use of a linear functional response: the effect of a mutualist on its partner’s per capita growth rate is linearly proportional to the mutualist’s density. Early models of obligate mutualism also shared this feature. Albrecht et al. (1974), May (1976), Christiansen and Fenchel (1977), and Vandermeer and Boucher (1978) introduced the idea of modeling mutualism through the intrinsic growth rate, shifting it from positive, in the case of facultative mutualism, to negative for obligate mutualism. Using linear functional responses, they generally found that, first, two obligate mutualists cannot stably coexist and, second, stable coexistence is possible if one species is obligate and the other is not, depending on the strength of the mutualism. These papers and others (e.g, Wolin, 1985; DeAngelis et al., 1986) further postulated that mutualistic functional responses are nonlinear, and thus attributed the unrealistic behavior of the Lotka-Volterra and similar models to their use of a linear functional response. Nonlinear functional responses were later explicitly modeled (e.g., Wright, 1989; Holland et al., 2002; Holland and DeAngelis, 2010; Revilla, 2015), confirming that nonlinear functional responses can indeed stabilize mutualistic populations.

Each of the aforementioned mutualism models, regardless of the functional response, assumes linear intraspecific density dependence; i.e., logistic within-species dynamics. However, nonlinear density dependence has been observed in controlled laboratory populations of organisms with simple life histories, such as *Daphnia* sp. and other Cladocera (Smith, 1963; Smith and Cooper, 1982) and *Drosophila* spp. (Ayala et al., 1973; Gilpin and Ayala, 1973; Pomerantz et al., 1980), and in long-term datasets on species with more complex life histories (Stubbs, 1977; Fowler, 1981; Sibly et al., 2005; Coulson et al., 2008). Models that relax the assumption of linear intraspecific density dependence have been proposed for single species (e.g., Richards, 1959; Schoener, 1973; Turchin, 2003; Sibly et al., 2005) and communities with two or more competitors (Ayala et al., 1973; Gilpin and Ayala, 1973; Schoener, 1976; Goh and Agnew, 1977; Gallagher et al., 1990), but never for mutualism (but see a generalized Verhulst-Lotka-Volterra model in Ribeiro et al. 2014 and a specific facultative-obligate model in Wang 2016). Given the prevalence of nonlinear intraspecific density dependence, and its known inuence on dynamics in other ecological contexts, the dearth of mutualism models that assume anything besides logistic growth suggests that our understanding of mutualistic dynamics may be quite incomplete.

In sum, the Lotka-Volterra mutualism model makes two separate assumptions that are likely violated in many natural systems: a linear effect of mutualistic interactions, and linear intraspecific density dependence. The former is widely thought responsible for the Lotka-Volterra mutualism model’s unrealistic behavior, but since the latter has never been investigated in the context of mutualisms, the relative importance of these two simplifying assumptions remains unclear. While we agree that many mutualistic interactions are likely nonlinear, the same could be said of competitive interactions, and yet Lotka-Volterra competition models endure. Is the need to eschew linear interaction rates truly fundamental for mutualisms? We approached this line of inquiry by returning to the original Lotka-Volterra mutualism model. To complement what is already known, we relax the assumption of linear intraspecific density dependence while leaving the assumption of a linear mutualistic functional response intact. We accomplish this by using the *θ*-logistic equation, which can decelerate or accelerate as a function of intraspecific density. We found that any accelerating model was always stable, and that decelerating models were stable with weak mutualism. We there-fore conclude that relaxing *either* of the Lotka-Volterra model’s major simplifying assumptions can prevent unrealistic model behavior. Given that nonlinear intraspecific density dependence appears to be widespread, nonlinearity in mutualistic interaction rates may be less important for stabilizing mutualisms than was previously believed.

## 2. Methods

The Lotka-Volterra mutualism model for population densities of two species, *N*_1_ and *N*_2_, takes the form

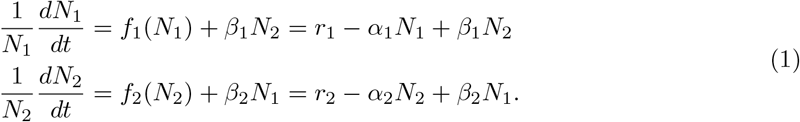

That is, the per capita change in population *i*’s density is a function of intraspecific density, *f*_*i*_ (*N*_*i*_), and a linear function of mutualist partner density, *β*_*i*_*N*_*j*_. It is further assumed that intraspecific density dependence, *f*_*i*_(*N*_*i*_), is logistic. This means the per capita growth rate approaches *r*_*i*_ when *N*_*i*_ approaches 0, and linearly decreases as intraspecific density increases, with slope −*α*_*i*_. Assuming positive parameter values, eq. (1) has the following behavior: each population grows when rare, each population has a stable positive abundance in the absence its mutualist partner, a feasible 2-species equilibrium exists if *β*_*i*_*β*_*j*_ < *α*_*i*_*α*_*j*_, and unbounded exponential growth occurs if *β*_*i*_*β*_*j*_ ≥ *α*_*i*_*α*_*j*_ (Vandermeer and Boucher, 1978).

The first terms in eq. (1) have not received the same scrutiny as the last terms. We suspect this has more to do with the ubiquity of the logistic model than careful evaluation of its application here. To explore this, we relax the assumption of logistic growth—the assumption that the difference between per capita births and deaths linearly decreases as density increases. We relax this assumption by modeling per capita growth rates using the *θ*-logistic model. This causes the per capita growth rate to be a decelerating function of density if the exponent (*θ*) is < 1 and an accelerating function if it is > 1 (Fig. 1). An exponent of 0 yields a density independent model and an exponent of 1 recovers the logistic model. We write each density dependent term, 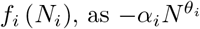:

**Figure 1:**
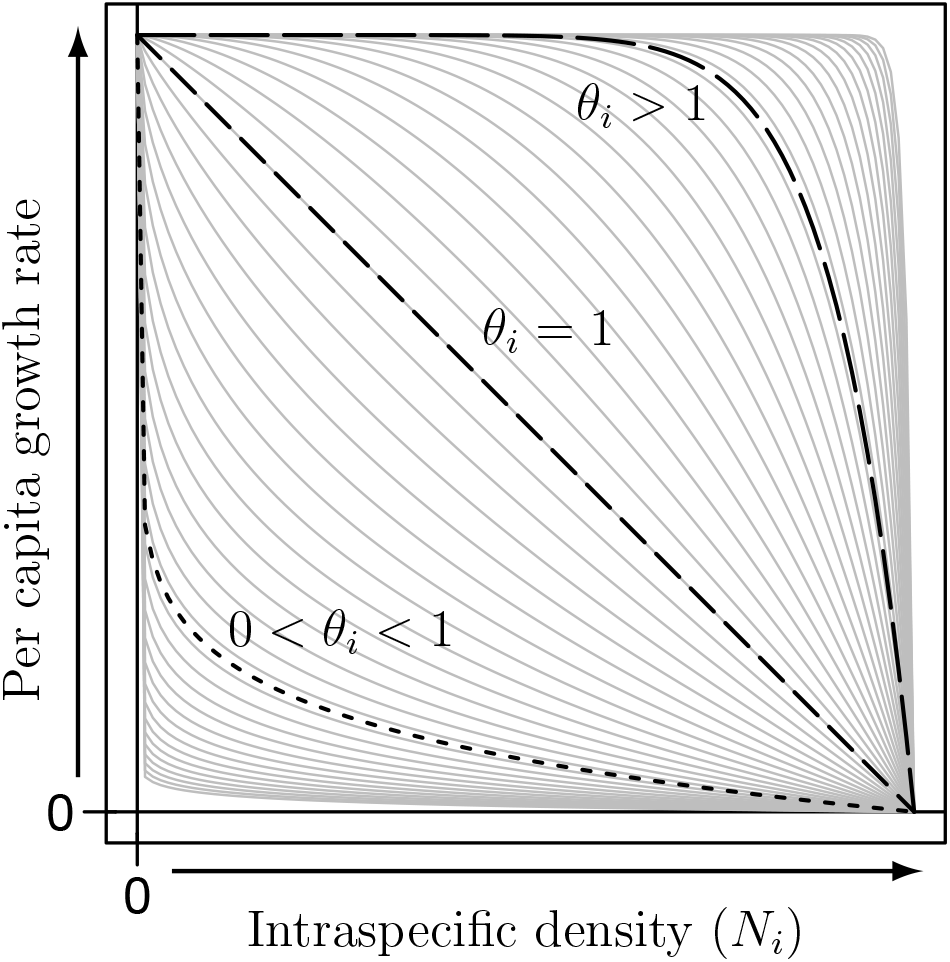
Values of *θ*_*i*_ used in eq. (2) to represent nonlinear per capita growth rates before accounting for the effects of mutualism. The figure shows how the per capita growth rates change as a function of intraspecific density, *N*_*i*_. The actual values used for numerical analyses are presented in light gray, with highlighted examples of decelerating intraspecific density dependence (*θ*_*i*_ = 1/10; short dashes, **.....**), linear intraspecific density dependence (*θ*_*i*_ = 1; medium dashes,**---**), and accelerating intraspecific density dependence (*θ*_*i*_ = 10; long dashes, **––**).

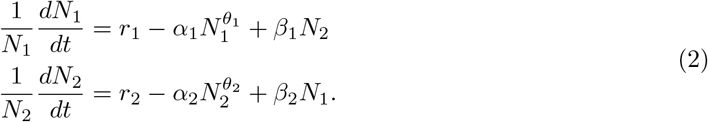

Our main experiment involved assessing stability of eq. (2) by modifying the four types of intraspecific density dependence (density independent, decelerating, linear, accelerating) in a model of mutualism with a linear functional response. Additionally, in the Supplementary Material, we (1) modeled per capita birth and death rates as separate nonlinear functions, each with their own exponent, (2) considered when exponents are different between the two populations, and (3) used a saturating functional response instead of a linear one using the procedures described in the remainder of 100 this section. A combination of analytical, numerical, and graphical techniques were used to assess the behavior of eq. (2). Specifically, we (i) found equilibria and (ii) determined the behavior around each equilibrium using local stability analysis. When analytical solutions were not possible (i.e., *θ*_*i*_ ≠ 0 or 1), we solved for stable equilibria numerically using the Livermore Solver for Ordinary Differential Equations, Automatic (LSODA) (Hindmarsh, 1983; Petzold, 1983) and solved for unstable equilibria using Newton’s method. LSODA is an integrator that was used because of its generality and ability to automatically handle stiff and non-stiff initial value problems, which were properties of our models. Newton’s method is an iterative root-finding algorithm we used to find unstable equilibria to a precision of 10^−15^, across state-space, from *N*_*i*_ = 0–10^100^ by orders of 10. Analyses were conducted in the R language and environment (R Core Team, 2016), with LSODA implemented in the deSolve package (Soetaert et al., 2010; Soetaert, 2010) and Newton’s method in the rootSolve package (Soetaert and Herman, 2009; Soetaert, 2009). Graphical analyses were conducted using a modified version of the R package phaseR (Grayling, 2014). Specifically, phase plots were created, using direction fields and zero-growth isoclines (i.e., nullclines) to corroborate and visualize our numerical findings. Code to run our analyses can be found at https://github.com/dispersing/Mutualism-NonlinearDensityDependence.

Parameter values for numerical analyses focused on the type of nonlinear per capita intraspecific density dependence (i.e., *θ*_*i*_) and the strength of mutualism (i.e., *β*_*i*_). For both of these types of parameters, we considered values ranging from 10^−2^–10^2^. The other parameter values (*r*_*i*_ and *α*_*i*_) did not qualitatively affect the model behavior in terms of number or stability of equilibria (C. Moore, unpublished results), so we do not discuss their effects in detail.

## 3. Results

*General results*. For all analyses with linear functional responses we found between 3 and 5 non-negative equilibrium population sizes (Fig. 2). Analytically, we found that (0,0) was always an equilibrium and always unstable. Further, there were always two boundary equilibria (*N*_1_ > 0, 0) and (0, *N*_2_ > 0), both of which were saddle nodes. The instability of the trivial and boundary equilibria means that populations always grow when rare, as expected. Numerically, we found that in cases where interior equilibria were present 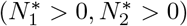, there were either one or two points. In cases where there was only one equilibrium point, it was always stable; in cases where there were two equilibrium points, the point proximal to the origin (0,0) was always stable and the point distal to the origin was a saddle node. Fig. 3 shows the six qualitatively different types of phase planes found in this study: (i) a trivial density independent case *θ*_*i*_ = 0; (ii & iii) unstable and stable configurations when intraspecific density dependence was decelerating, 0 < *θ*_*i*_ < 1; (iv & v) unstable and stable configurations when intraspecific density dependence was linear, *θ*_*i*_ = 1; and (vi) a stable configuration when intraspecific density dependence was accelerating, *θ*_*i*_ > 1.

**Figure 2:**
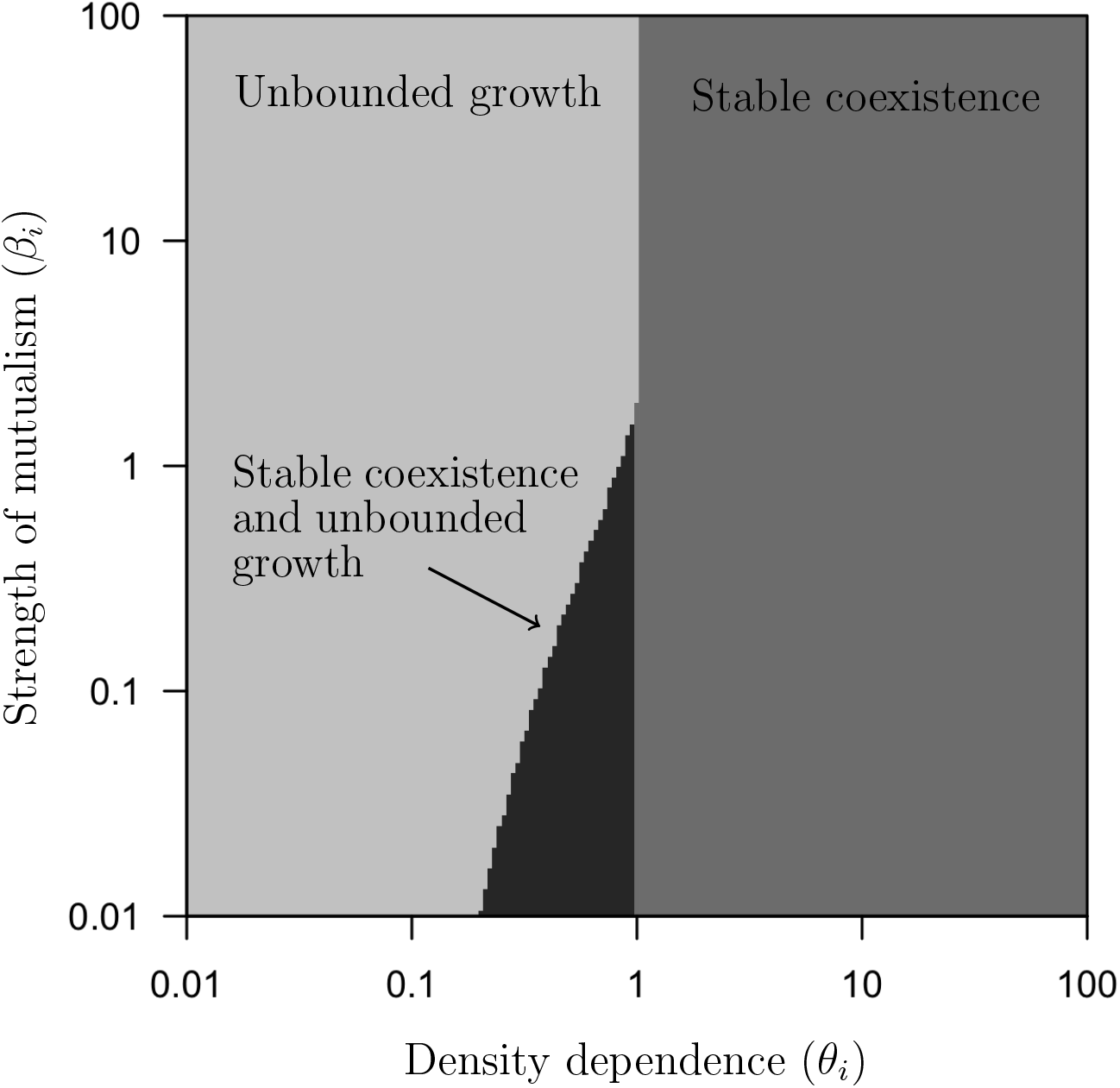
Number of equilibrium points (shades of gray) across all values of intraspecific density dependence (*θ*_*i*_) and strength of mutualism (*β*_*i*_), while holding the remaining parameters constant at *r*_*i*_ = 4, *α*_*i*_ = 2. Across all analyses, there were always between 1 and 2 interior equilibria (3 and 5 total equilibria, including the trivial and boundary equilibria). The light-gray regions corresponds to unstable configurations where no interior equilibrium existed, the medium-gray regions correspond to stable configurations where one stable interior equilibrium existed, and the dark-gray regions correspond to areas with two interior equilibria, one stable at low densities and one saddle at high densities.

**Figure 3:**
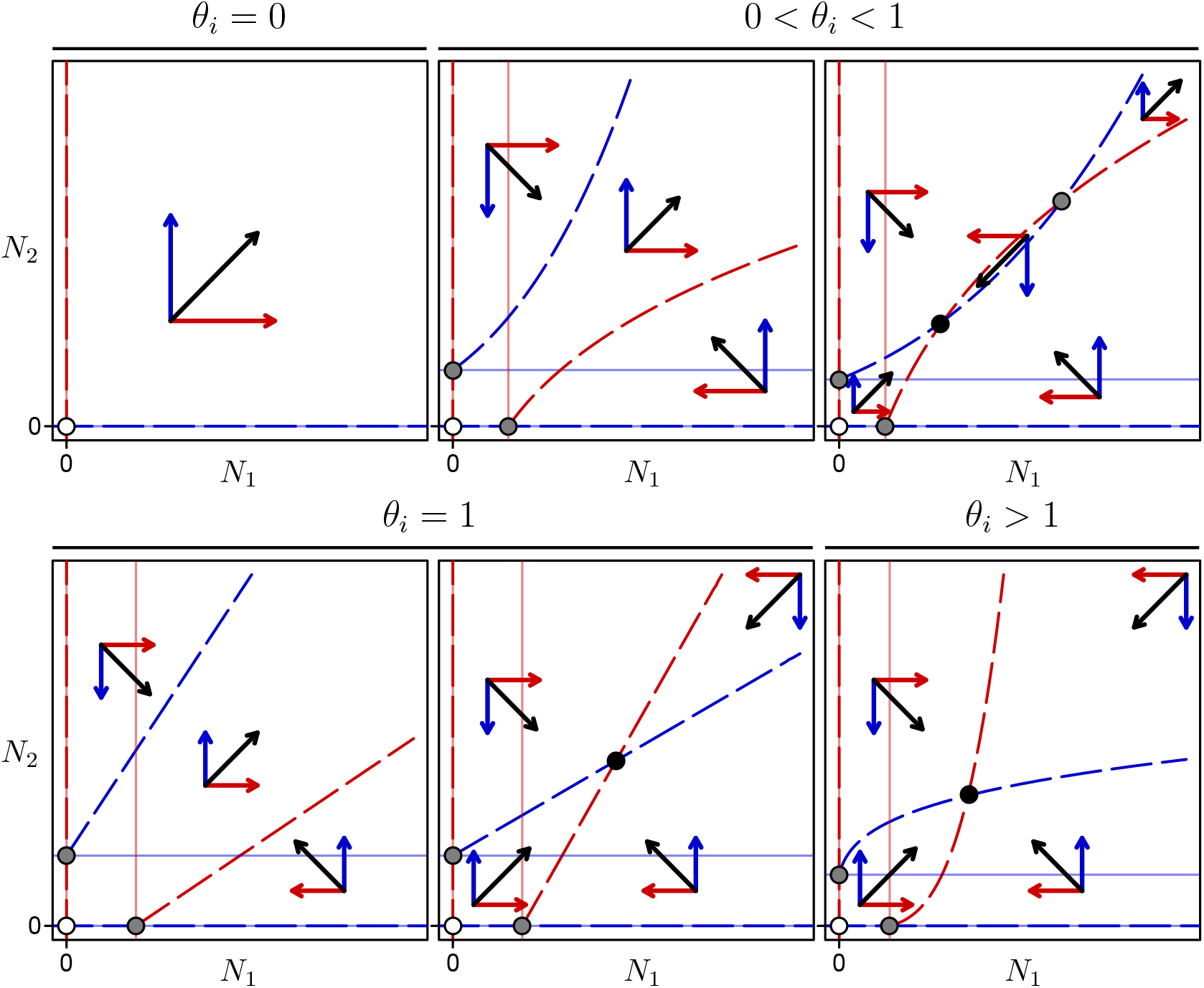
Phase planes representing the qualitative dynamics of 2-species mutualistic interactions for different models of per capita intraspecific density dependence. Each panel shows the densities of *N*_1_ and *N*_2_ on the *x*- and *y*-axes. Within each panel, zero-growth isoclines (nullclines) are shown for *N*_1_ (red) and *N*_2_ (blue): (i) when there is no mutualism (*β*_*i*_ = 0) as solid, light lines (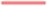 or 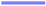) and (ii) when mutualism is present (*β*_*i*_ > 0) as dashed lines (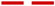 or 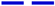). Arrows within panels show the qualitative direction vectors for *N*_1_ (red), *N*_2_ (blue), and together (black) for all changes in direction for each phase plane. Points within panels represent unstable (white), stable (black), or saddle nodes (gray). The trivial intraspecific density independent result (*θ*_*i*_ = 0) is shown in the *top-left* panel, the two results of decelerating intraspecific density dependence (0 < *θ*_*i*_ < 1) are shown in the *top-center and -right* panels, linear intraspecific density dependence (*θ*_*i*_ = 1) is shown in the *bottom-left* and -*center* panels, and accelerating intraspecific density dependence (*θ*i > 1) is shown in the *bottom-right* panel.

In general, in the absence of mutualism, decelerating intraspecific density dependence increased both species’ densities at equilibrium (*β*_*i*_ = 0 plane in Fig. 4, left panel). Oppositely, accelerating intraspecific density dependence decreased the equilibrium densities. Strong mutualism (high *β*_*i*_) destabilized populations with decelerating intraspecific density dependence, but populations with accelerating intraspecific density dependence were always stable (Fig. 4, center panel; note that only stable equilibria are shown, so missing portions of the surface at high *β*_*i*_ and low *θ*_*i*_ denote loss of stability; see also Supplemental Material, section 2). Further, when a stable interior equilibrium was present, adding mutualism to populations with decelerating intraspecific density dependence generated a larger benefit of mutualism than with accelerating intraspecific density dependence (Fig. 4, right panel).

**Figure 4:**
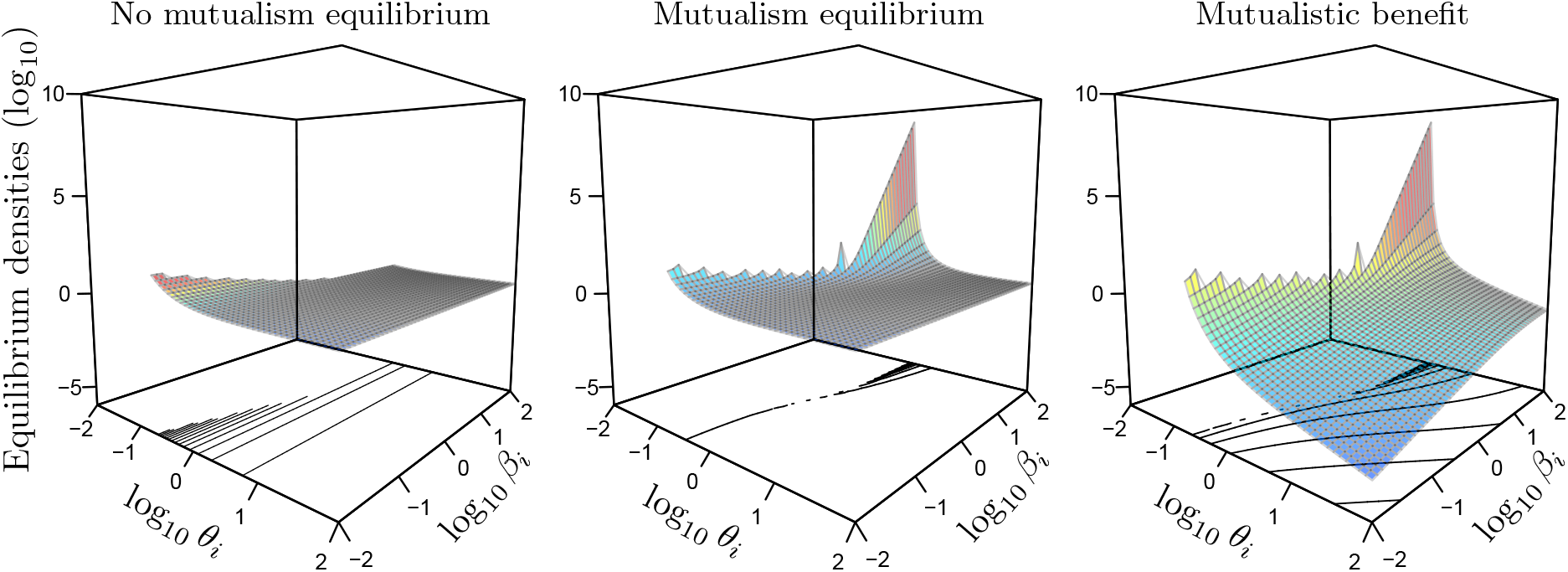
For model (2), nonlinear per capita growth rates with a linear functional response of mutualism, the location of the interior equilibrium in the absence of mutualism (*β*_*i*_ = 0, left), stable interior equilibrium with mutualism (center), and the benefit of mutualism as the difference between the two (right). The locations of equilibria were identified as the Euclidean distance from the origin, 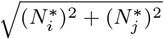, for identical parameters for each species: *r*_*i*_ = 4, *α*_*i*_ = 2. Each panel shows the aforementioned response on the vertical axis, the type of intraspecific density dependence (*θ*_*i*_ from 10^−2^–10^2^) on the left horizontal axis, and the strength of mutualism (*β*_*i*_ from 10^−2^–10^2^) on the right horizontal axis. Additionally, each panel shows the relative values of each surface (colors), the absolute values of each surface (same axes across panels), and contour lines at the base of each plot show changes in the surface. Further, in areas where there is no surface, there was no stable interior equilibrium when *β*_*i*_ ≠ 0 (center). In the left panel without mutualism, there were stable interior equilibria across all values of *θ*_*i*_, but we removed the same part of the surface to aid comparison across panels.

*Decelerating density dependence*, 0 < *θ*_*i*_ < 1. When 0 < *θ*_*i*_ < 1, we found that there were 1–2 interior equilibria (3–5 total equilibria), depending on the strength of mutualism. In the absence of mutualism, the interior equilibrium (and consequently the boundary equilibria by setting either coordinate to 0) is at

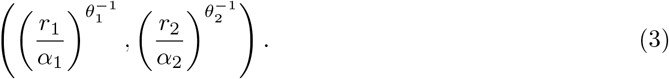

Notice the 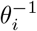 exponent. In these cases of decelerating density dependence, as *θ*_*i*_ decreases from 1, the greatest change in growth rate occurs at lower densities (Fig. 1). Furthermore, the equilibrium density in the absence of mutualism grows larger as *θ*_*i*_ decreases.

Adding mutualism to populations with decelerating density dependence changed the dynamics in either of two ways: (i) it destabilized the populations resulting in unbounded population growth (Fig. 3, top-center panel) or (ii) it created both a stable and saddle node (Fig. 3, top-right panel). For very small values of *θ*_*i*_, populations were always unstable with mutualism added (i.e., *β*_*i*_ > 0). As decelerating density dependence became more linear (i.e., as *θ*_*i*_ → 1), however, weak mutualism (small values of *β*_*i*_) resulted in an alternative configuration in which zero-growth isoclines crossed twice. Of these two equilibria, the stable equilibrium point was always larger than in the absence of mutualism (*β*_*i*_ = 0) and the saddle node was always larger than the stable point. For the same values of *θ*_*i*_ with stable and saddle nodes, increasing *β*_*i*_ increased the stable point and decreased the saddle point. Continuing to increase *β*_*i*_ ultimately resulted in a saddle-node bifurcation, beyond which all configurations were unstable, illustrated as the light-dark gray boundary in Fig. 2.

*Linear density dependence*, *θ*_*i*_ = 1. When *θ*_*i*_ = 1, there were either 0 or 1 interior equilibrium configurations (3 or 4 total equilibria) that respectively corresponded to the absence of presence of an interior stable point. Linear density dependence is equivalent to the most traditional formulation of mutualism, the Lotka-Volterra competition model with the sign reversed of the effect of another population. Although the behavior of this model is well-known, we summarize its properties briefly here for ease of comparison. In the absence of mutualism, the interior equilibrium (and consequently the boundary equilibria by setting either value to 0) is at

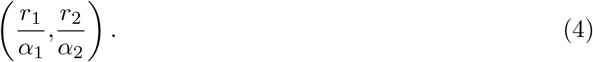

The slope of the zero-growth isocline as it increases from the boundary equilibrium is 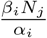, and zero-growth isoclines form a stable interior equilibrium point anytime *β*_*i*_*β*_*j*_ < *α*_*i*_*α*_*j*_. This is equivalent to the more traditional notation, *α*_*i*_*α*_*j*_ < *α*_*ii*_*α*_*jj*_ found in ecology texts (e.g., May, 1981; DeAngelis et al., 1986; Kot, 2001). The location of the stable interior equilibrium point is

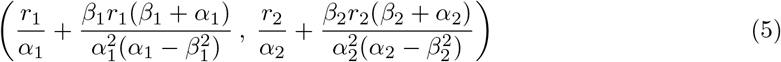

*Accelerating density dependence*, *θ*_*i*_ > 1. When *θ*_*i*_ > 1, there was always one interior equilibrium (4 total equilibria), irrespective of the strength of mutualism (Figs. 2, 4). In the absence of mutualism, the interior equilibrium is again given by (3). Again, note the 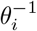 in the exponent. In these cases of accelerating density dependence, as *θ*_*i*_ increase from 1, the greatest change in growth rate occurs at higher densities (Fig. 1). Furthermore, the equilibrium point in the absence of mutualism decreases as *θ*_*i*_ increased (Fig. 4, left panel). With mutualism (*β*_*i*_ > 0), in addition to always being stable, the benefit decreased as *θ*_*i*_ increased.

*Supplementary Material: Births and deaths as separate processes, interspecific differences in intraspecific density dependence, and saturating functional response*. Assuming per capita birth and death rates were independent processes, we modeled them as separate nonlinear functions. Our main finding was that as long as one of the exponents was accelerating, the interior equilibrium point would always be stable. We found no qualitatively new model behaviors when we allowed the two species to have differently shaped intraspecific density dependent functions. Notably, the presence of 187 a single, stable interior equilibrium point is possible as long as *either* species has accelerating density dependence. We also replaced the linear functional response in eq. (2) with a saturating functional response, finding that the saturating function would always stabilize the interior equilibrium point, but with less mutualistic benefit (the difference in density in the absence and presence of a mutualist partner).

## 4. Discussion

Lotka-Volterra models of mutualism assume that intraspecific density linearly decreases per capita growth rates. Other population models of mutualism have inherited this assumption and have generally concluded that 2-species models of mutualism are inherently unstable. In real populations, however, not only do nonlinear per capita growth rates exist, but they seem to be the rule rather than the exception (Stubbs, 1977; Fowler, 1981; Sibly et al., 2005). In this study, we examined how relaxing the assumption of linearly dependent per capita growth rates affected stability and mutualistic benefit in these models. We found that when per capita growth rates decrease most strongly at low densities and are decelerating, mutualism usually destabilizes the model. In contrast, when growth rates decrease most strongly at high densities and are accelerating, models are always stable with mutualism. Despite the tendency for mutualism to destabilize the 2-species equilibrium with decelerating density dependence, the benefit was greater compared to stabilizing, accelerating density dependence.

Our paper presents an alternative way that the classic Lotka-Volterra mutualism model can be modified to stabilize mutualism. Simply put, we added a layer of biological realism (nonlinear intraspecific density dependence) to the Lotka-Volterra mutualism model and we found informative ways that within-species properties could stabilize mutualism, even with a linear functional response modeling the interaction between species. Support for decelerating and accelerating density dependence has largely been based on large datasets from observational studies (e.g., 1750 species of mammals, birds, fish, and insects in Sibly et al., 2005). Further work to determine whether species that engage in mutualism are more likely to have accelerating intraspecific density dependence, which we found to be stabilizing, would be useful.

From an ecological perspective, species’ nonlinear responses to intraspecific density arise from differences in ecological habits or population structure. Sedentary organisms, like many plants for example, exhibit a more-or-less-constant death rate at low-to-intermediate population densities, and then at higher densities death rates tend to rapidly increase (as in scramble competition or self-thinning, Yoda et al., 1963) or increase linearly (as in contest competition, Crawley and Ross, 1990), resulting in accelerating density dependence. Subsets of populations, such as age or stage, can experience different vital rates and generate nonlinear density dependence for populations as a whole. In African ungulates, for example, increases in density led to increases in adult mortality, while juvenile mortality remained relatively constant at all population densities (Owen-Smith, 2006). In fact, many mutualisms occur between species with structured populations, so our study may lend insights into these interactions. As examples, many plant-mycorrhizal associations are mutualistic in the seedling stage (Grime et al., 1987; Jones and Smith, 2004; van der Heijden and Horton, 2009)and adult plants engage in mutualistic interactions with pollinators when they reach a reproductive stage.

From an evolutionary perspective, a long-standing theory about why we see nonlinear density dependence comes from evolutionary theories of life-history strategies; i.e., *r*- and *K*-selected species (Gilpin and Ayala, 1973; Stubbs, 1977; Fowler, 1981), including *θ*-selection (Gilpin et al., 1976). Setting aside historical controversies, this body of theory has generated very useful quantities like the specific growth rate of a population. The most general prediction made is that populations with a high specific growth rate (commonly referred to as *r*-selected) should exhibit decelerating density dependence since their survival probability drops off precipitously at relatively low densities. On the other hand, populations with a low specific growth rate (commonly referred to as *K*-selected) should exhibit accelerating density dependence since their survival probabilities drop off at relatively high densities (see Figs. 1, 2 in Fowler, 1981). Based on our study we suspect that different life-history strategies may both be a result of and a causative factor in the evolution of mutualistic interactions, and further work should examine how engaging in a mutualistic interaction should change the shape of density dependence and how changing density dependence affects a species ability to engage in a mutualistic interaction.

In conclusion, the linear functional response has historically been the scapegoat for theoretical studies of the population dynamics of mutualism. For example, the eminent Lord Robert May (1976) writes:

> *… the simple, quadratically nonlinear, Lotka-Volterra models ... are inadequate for even a first discussion of mutualism, as they tend to lead to silly solutions in which both populations undergo unbounded exponential growth, in an orgy of mutual benefaction. Minimally realistic models for two mutualists must allow for saturation in the magnitude of at least one of the reciprocal benefits.*

In this paper, we build on May’s idea of modifying the Lotka-Volterra mutualism model; not through the saturation of benefits, but through intraspecific density dependence. We found that biologically-realistic nonlinear density dependence significantly changes the dynamics of the original Lotka-Volterra mutualism model, where we found that accelerating density dependence always stabilized our models but with weaker mutualistic benefit relative to decelerating density dependence. We hope that this study will further stimulate ecologists to consider all simplifying of assumptions of even the most basic models and also to investigate more deeply into the relationships between intraspecific density, interspecific density, and population growth to gain a better grasp on mutualistic population dynamics.

## 5. Acknowledgements

We thank Katie Dixon, Frances Ji, Brian Lerch, Robin Snyder, and Chris Steiha for comments on an early draft of the manuscript. K.C.A. and C.M.M. were supported in part by a James S. McDonnell Foundation Complex Systems Scholar award to K.C.A.

